# Microhomology-Driven Genomic Alterations in Cancer Genomes: Patterns, Prevalence, and Clinical Implications

**DOI:** 10.64898/2026.09.03.744669

**Authors:** Daria Kostka, Paweł Sztromwasser, Marek Kimmel, Roman Jaksik

## Abstract

Homologous recombination deficiency (HRD) can force cancer cells to rely on alternative DNA repair pathways, including microhomology-mediated end joining (MMEJ), an error-prone mechanism that can generate deletions with microhomology at repair junctions. Because such patterns may reflect DNA repair defects with clinical relevance, this study aimed to assess the biological and prognostic significance of microhomology-associated deletions and optimize their detection parameters to improve patient stratification and inform targeted therapeutic strategies, including PARP inhibition.

We optimized the microhomology length threshold (M_lt_ parameter) using signal-to-noise ratios and survival models. We evaluated the utility of whole-exome (WES) versus whole-genome sequencing (WGS). We then investigated how Loss of Function (LoF) alterations affect microhomology-associated deletion burden and assessed their prognostic significance across an ovarian cancer (OV) cohort and a TCGA Pan-Cancer dataset.

Applying an M_lt_ = 2 threshold maximized the precision and statistical reliability of microhomology-associated deletion identification. WGS yielded higher reproducibility, whereas WES proved insufficient for accurate estimation. Inactivation of tumor suppressor genes, including RB1 and CCDC122, was associated with increased microhomology-associated deletion burden. Importantly, higher burden correlated with extended overall survival in ovarian cancer. Furthermore, while baseline microhomology-associated deletion burden varies by tumor type, Pan-Cancer analysis identified candidate gene-level alterations, including FDX1 and PDE8B, whose LoF increased microhomology-associated deletion burden across tumors.

Precise parameterization combined with WGS resolution highlights the extent to which specific gene losses affect deletion burden consistent with MMEJ-mediated repair. Identifying these genetic vulnerabilities and the resulting deletion burden may provide a candidate prognostic biomarker and foundation for personalized therapies.

## Introduction

DNA double-strand breaks (DSBs) are among the most severe forms of genomic damage, with the potential to cause mutations, genomic instability, and cancer onset if not properly repaired (1). Cells address DSBs through several repair mechanisms, including homologous recombination (HR) and nonhomologous end joining (NHEJ). HR relies on a homologous sequence as a template, ensuring high-fidelity repair by using sister chromatids during the S and G2 phases of the cell cycle (2,3). In contrast, NHEJ operates without a template and can be further categorized into canonical NHEJ (C-NHEJ) and alternative NHEJ (alt-NHEJ), also known as microhomology-mediated end joining (MMEJ) (4). While both NHEJ and MMEJ can result in small deletions at the break site, MMEJ is distinct in that it relies on microhomologous sequences flanking the break. This pathway is particularly error-prone, leading to genomic rearrangements, translocations, and deletions. MMEJ has been shown to play a critical role in HR-deficient tumors (3,5).

MMEJ proceeds in multiple steps, involving DNA end resection, annealing of microhomologous sequences, removal of non-complementary regions, gap filling via DNA synthesis, and ligation to restore strand continuity (6). A key similarity between MMEJ and HR is the shared initial step of DSB end resection, which leads to the formation of 3’ single-stranded DNA (ssDNA) overhangs coated with replication protein A (RPA) (2). When the BRCA1 and BRCA2 genes are mutated in a cell, the cell directs DSBs to the MMEJ repair pathway. However, the error-prone nature of MMEJ can lead to the loss of genetic information at the break site and may result in translocations and complex rearrangements (7).

The MMEJ mechanism begins with PARP-1 competing with Ku70/80 for binding to DSB ends, recruiting MRN and CtIP to initiate resection. This process generates 3’ ssDNA through endonucleolytic cleavage followed by exonucleolytic digestion. HMCES binds to ssDNA to protect the free ends (8,9). End bridging and microhomology alignment are mediated by PARP-1, MRN, and polymerase theta (Polθ). Next, non-homologous flaps are removed: 3’ tails by ERCC1/XPF or FEN1, and 5’ tails by FEN1. Polθ fills the gaps, uniquely initiating synthesis and frequently causing templated insertions. The process concludes with ligation by LIG3/XRCC1 (8,10).

Homologous recombination deficient (HRD) cells with mutations in BRCA1/2 utilize MMEJ as a backup repair pathway (11). In such cases, Polθ inhibitors are emerging as a novel therapeutic option for BRCA1/2-mutated tumors and in cases of PARP inhibitors (PARPi) resistance (12). RB-dependent tumors, which also rely on MMEJ, are highly sensitive to PARPi such as olaparib, and their therapeutic efficacy is enhanced when combined with etoposide, offering good outcomes in retinoblastoma treatment. RB inactivation, observed in tumors such as melanoma, small cell lung cancer, and triple-negative breast cancer, significantly increases sensitivity to PARPi, often surpassing the response seen with BRCA mutations (12). Furthermore, Polθ is overexpressed in HRD-associated cancers, including breast, lung, bladder, colorectal, gastric, glioma, pancreatic, prostate, melanoma, and uterine cancers, potentially linked to the underlying HR deficiency (13).

Ovarian cancer, characterized by HRD, relies on alternative DNA repair mechanisms such as microhomology-mediated end joining (MMEJ). The high mortality associated with this cancer primarily stems from late diagnosis, with up to 75% of cases identified at an advanced stage (14). Its non-specific early symptoms hinder early detection (15). Molecularly targeted therapies offer a promising alternative or a complement to traditional chemotherapy in the treatment of ovarian cancer. Currently, bevacizumab and olaparib are approved by the U.S. Food and Drug Administration (FDA), while rucaparib has received accelerated approval for the treatment of advanced BRCA-mutated ovarian cancer following at least two lines of chemotherapy (14). Furthermore, numerous clinical trials investigating novel PARPi have the potential to improve outcomes. Ultimately, overcoming platinum resistance and recurrent ovarian cancer requires innovative strategies and therapies, rigorously tested in clinical trials (14).

The MMEJ repair mechanism, which utilizes microhomologous sequences for end joining, can be accompanied by template switching during DNA replication. According to the fork stalling and template switching (FoSTeS) model, stalled replication forks can switch templates, aligning with complementary microhomology, allowing DNA synthesis to continue (16). This process can lead to structural rearrangements such as deletions, insertions, and multi-nucleotide variants (MNVs). Template switching can occur over considerable genomic distances, spanning hundreds of kilobases or even megabases and thereby contributing to extensive genomic alterations (17). Furthermore, repetitive elements and secondary DNA structures can induce DSBs, additionally promoting genomic instability. In such cases, microhomology plays a dual role, facilitating both template switching and initiating replication at a new fork, potentially leading to complex rearrangements (17,18).

In addition to the characteristic microhomology observed at deletion junction sites, MMEJ is also associated with the formation of small insertions. These insertions arise through DNA synthesis, frequently utilizing the flanking sequences around the DSB as a template. The presence of polymerase theta (Polθ) plays a crucial role in this process, as it extends short microhomologies (≥1 bp) at the DSB ends, enabling gap filling and facilitating repair (19). In some cases, the repair process may involve template switching prior to ligation, leading to templated insertions. This occurs when the extended intermediate repair product dissociates and re-anneals at a different microhomology site, incorporating additional nucleotides into the final repair product. Such template switching events can lead to insertions of varying degrees of complexity (20).

The primary aim of our research was to identify characteristic features of tumors, which led to the development of an algorithm enabling the classification of indels (insertions, deletions) and MNVs as potentially dependent on MMEJ repair mechanisms. To implement the detection algorithm, we developed a custom tool within the R environment. While pre-existing software packages such as Del-read (21), MHcut (22), MMEJ_detection (23), and YAPSA (24) address this task, our tailored implementation provided greater flexibility for parameter optimization and enabled the expansion of our analysis to incorporate template switching. Specifically, this enabled us to optimize the microhomology length threshold (*M_lt_*) parameter to increase the precision of microhomology-associated deletion classification. Additionally, a comparative analysis of WGS and WES data for microhomology detection was performed, allowing us to assess the impact of inherent WES limitations on the ability to identify potential MMEJ signatures. A key element of the research also involved assessing the relationship between LoF and microhomology-dependent deletion burden, enabling us to determine the extent to which inactivation of selected genes likely promotes cell switching to alternative DNA repair pathways. This tool was designed to analyze genomic features associated with HRD and its impact on tumor biology. By providing a precise method for identifying microhomology-associated alterations, our algorithm deepens the understanding of MMEJ activity in HRD-dependent tumors. The findings presented in this publication will be crucial for better patient stratification and optimizing the use of targeted therapies, such as PARPi, ultimately contributing to more effective treatments with reduced toxicity and improved clinical outcomes.

## Data

Genomic data derived from The Cancer Genome Atlas (TCGA) and the International Cancer Genome Consortium (ICGC) were utilized in this study, with a specific focus on ovarian cancer (OV) and a range of other cancer types. The dataset includes somatic variants detected using both Whole Genome Sequencing (WGS) and Whole Exome Sequencing (WES).

### Ovarian Cancer Data

The ovarian cancer cohort included:

- 121 patients analyzed using the WGS method.
- 506 patients analyzed using the WES method.

Somatic mutations in WGS data were identified using Strelka2 (v2.9.10) (25) in matched tumor-normal mode. WES data were processed using: MuTect2 (v4.2.4.1) (26), Pindel (27) (v0.6.3) and VarScan2 (v6.0) (28). Functional annotation of variants was performed using Ensembl VEP (v102) (29).

A common subset of 23 samples from ovarian cancer patients was analyzed using both WGS and WES methodologies. This overlap enabled comparative evaluation of the two sequencing approaches within the same cohort.

### Pan-cancer dataset

To further investigate microhomology-mediated deletions, we analyzed WES data - 9,829 samples and WGS data - 8,393 samples across 32 cancer types.

Clinical metadata, current as of March 23, 2026, included patient information such as age, sex, disease stage, and treatment. The source of these data was the TCGA Clinical Data Resource (CDR) Outcome dataset (TCGA-CDR-SupplementalTableS1.xlsx) available on the GDC Data Portal (30).

## Methodology

We developed MHDetect, an R-based algorithm designed to detect and classify deletions, insertions, and MNVs associated with MMEJ repair. The tool parses input data directly from variant call format (VCF) files utilizing the VariantAnnotation package (31). The analysis is governed by four user-defined parameters: *k*, *M_lt_*, *genome*, and *Interval*. The *k* parameter defines the length of the breakpoint-adjacent sequence extracted upstream and downstream of each variant. In contrast, *M_lt_*, the minimum microhomology length threshold, defines the minimum number of consecutive matching nucleotides required at the variant junction to classify an event as microhomology-associated. Thus, *M_lt_* does not determine the size of the flanking window; it is a threshold applied after sequence comparison. The *genome* parameter specifies the reference genome assembly, and *Interval* determines the length of the flanking regions around an MNV utilized to simulate the template switching mechanism.

The MHDetect algorithm (Figure 1A) comprises distinct modules dedicated to analyzing deletions, insertions, and MNVs, with the primary objective of determining whether a given mutation is dependent on microhomology-related DNA repair. In the common preprocessing stage, the algorithm filters out the low-confidence variants. Subsequently, the input data are stratified into deletions, insertions, and MNVs, with each category assigned to a separate data frame: DEL, INS, and MNV. This preprocessing step is common for all downstream analyses.

**Figure 1.**
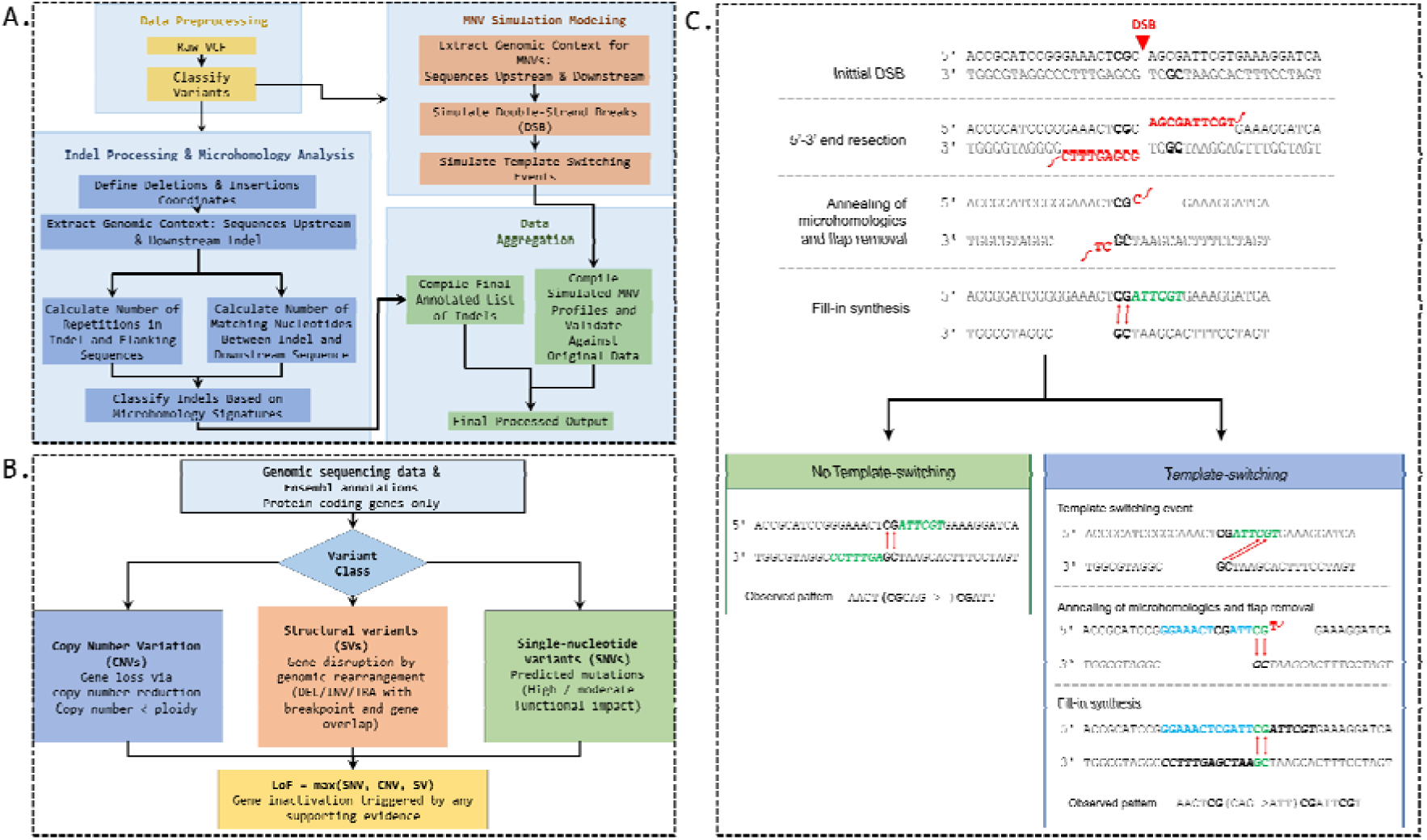
Overview of the MHDetect workflow, Loss of Function assessment, and DSB repair models. A. Schematic representation of the MHDetect algorithm workflow for identifying and classifying deletions, insertions, and multi-nucleotide variants (MNVs), including the simulation of template-switching events. B. Integrative algorithm for the assessment of Loss of Function (LoF) events, combining data from single nucleotide variants (SNV), short indels, copy number variants (CNV), and structural variants (SV). C. Mechanistic models of microhomology-mediated double-strand break (DSB) repair, comparing the standard pathway (No Template-switching, left) with complex genomic alterations generated via the Template-switching pathway (right).

### Deletions and Insertions

The module analyzing deletions and insertions follows a standardized workflow. Initially, the algorithm identifies sequences altered by deletions and compares them against the reference genome (Figure 1A). For each event, MHDetect extracts breakpoint-adjacent sequences upstream and downstream of the variant, with the length of each flank defined by the k parameter. In our analysis, k was set to 25 bp. This value was selected to cover the typical range of microhomology tracts reported for MMEJ, which are commonly described as short homologous sequences of approximately 2–25 bp flanking double-strand break junctions (32–34). Thus, k = 25 provides sufficient local sequence context to detect such microhomologies while limiting the search space to the immediate breakpoint neighborhood. An analogous extraction and comparison process is performed for insertions (Figure 1A).

Upon completing the individual evaluation of deletions and insertions, the results are merged into a single consolidated table for downstream analysis. The algorithm assesses the presence of mutation sequence repeats in the regions immediately following the mutation, as well as the number of consecutive matching nucleotides between the mutation and the following sequence. Based on these metrics, the indel is classified as either microhomology-mediated or repeat-mediated. Both deletions and insertions are classified as microhomology-mediated if the number of matching nucleotides is greater than or equal to the threshold value *M_lt_*. For our study, we set *M_lt_* = 2, which yielded optimal performance during parameter tuning.

### MNV

The analysis of MMEJ-associated MNVs involves simulating DSBs within a defined genomic region surrounding the MNV to reconstruct the microhomology-mediated DNA repair process, including both simple and template-switching pathways (Figure 1C). First, the algorithm extracts the flanking DNA sequences based on the specified length parameter. Subsequently, it simulates the formation of DSBs at every possible position within a predefined nucleotide range. For each simulated break, the adjacent DNA sequences on both sides are evaluated to determine the repair direction and identify potential microhomologies (Figure 1A).

Drawing on the template switching mechanism, the simulation algorithm generates potential DSB positions, calculates the center of the defined range, and extends it by a specified number of nucleotides to ensure all relevant sequence fragments are included in the simulation. The reconstructed repair process encompasses a sequence of operations, including the excision of non-homologous overhangs (flaps), the synthesis of missing nucleotides, and the rejoining of DNA strands guided by newly identified microhomologies. All potential repair trajectories are evaluated, and the optimal microhomology matches are recorded, enabling the precise calculation of deleted and inserted DNA fragment coordinates (Figure 1A). Finally, by comparing the positions and sequence composition of the simulated MNVs with the original observed MNVs, their similarity is evaluated to assess whether the observed event is consistent with a microhomology-associated repair model.

The algorithm returns comprehensive results detailing deletions, insertions, and MNVs, alongside their association with MMEJ repair. Specifically, it outputs distinct data frames containing the processed deletions and insertions, an aggregated summary of MNVs correlated with the simulation results, and the exhaustive outputs of the template switching simulations.

### Loss of Function Analysis

The identification of LoF events was based on the integration of four types genomic variants, including single-nucleotide variants (SNVs), short indels, structural variants (SVs), and copy number variations (CNVs). Gene annotations mapped to the GRCh38 reference genome were retrieved from the Ensembl database (release 115) (35). The extracted data encompassed gene identifiers, chromosomal locations, and genomic ranges, with the analysis restricted to protein-coding genes on chromosomes 1-22, X, and Y.

As illustrated in Figure 1B, the implemented algorithm evaluates gene inactivation through a modular framework that consolidates evidence from four distinct data streams. For SNVs and short indels, the mutation’s impact on protein function was categorized using Ensembl VEP (v115.2) impact scores, where variants classified as HIGH or MODERATE were designated as LoF events. CNV analysis assessed gene dosage relative to sample ploidy, identifying significant loss when the copy number fell below the rounded ploidy level. Structural variants were evaluated by mapping their genomic breakpoints to specific gene regions; deletions were marked as LoF if they overlapped with any portion of the gene, whereas duplications, inversions, and translocations were identified as LoF events if they disrupted the gene’s structural integrity-defined by one breakpoint located within the gene boundaries and the other externally. The final binary LoF status, where 1 signifies Loss of Function and 0 signifies retained function, was assigned based on a maximum-score integration, where a single positive criterion across any of the four sources was sufficient to classify a gene as at least partially inactivated.

Because our definition integrates heterogeneous variant classes, including moderate-impact SNVs and copy-number losses, LoF status should be interpreted as a putative functional alteration rather than definitive biallelic gene inactivation.

### Statistical analysis and data visualization

All statistical analyses and visualizations were performed in the R environment (version 4.5.3). The following packages were used: ggplot2, ggpubr, plotly, ggrepel, cowplot, smplot2, corrplot, survival, survminer, aod, betareg, MASS, and survRM2. Statistical significance was defined as a two-sided p-value < 0.05. To account for multiple testing, p-values were adjusted using the Benjamini-Hochberg false discovery rate (FDR) correction where applicable.

### Optimization of microhomology length threshold and stability analysis

To determine the optimal *M_lt_* for reliable detection of deletions associated with the MMEJ mechanism, a signal-to-noise ratio (SNR) analysis was performed. Expected counts of microhomology-associated events were estimated using a shifted genomic background window (+100 nt) and compared with observed counts derived from regions immediately downstream of indels. The prognostic impact of the optimized parameter *M_lt_* was further evaluated using hazard ratios (HR) and the concordance index (C-index) in survival models across different threshold settings.

The stability of LoF-gene associations was assessed using downsampling approaches. For each classification parameter *M_lt_* (ranging from 1 to 11), deletions classified as microhomology-mediated at threshold *M_lt_* were randomly downsampled to match the number of samples classified at threshold *M_lt_* + 1. A total of 100 bootstrap iterations were performed for each value of *M_lt_* to evaluate the distribution of significant genes and the robustness of effect size estimates (median log_2_ fold change). The non-downsampled deletions for both *M_lt_* and *M_lt_* + 1 were retained as reference points for comparison.

To identify potential systematic biases and validate statistical thresholds, quantile-quantile (QQ) plots were generated by comparing observed and expected -log_10_(p-values), using the median p-value obtained across the 100 bootstrap iterations for each gene. The reproducibility of gene-level associations was further quantified using empirical cumulative distribution functions (CDFs), representing the proportion of genes remaining significant across repeated downsampling iterations.

### Mutation profiling and variant caller concordance assessment

Basic statistical profiling included quantification of total counts and proportions of deletions associated with microhomology and tandem repeats. The distribution of deletion classes (microhomology-mediated, repeat-mediated, and others) was compared across variant calling algorithms (VarScan2 WES, Pindel WES, MuTect2 WES, MuTect2 WGS, and Strelka2 WGS). Differences in the estimated contribution of microhomology-associated deletions between WES and WGS pipelines were visualized using stacked bar plots and boxplots.

A Spearman correlation matrix (correlogram) was constructed to assess consistency of microhomology-associated deletion proportions estimated from WES-based algorithms. Cross-platform concordance was additionally evaluated by comparing median overall indel burdens between matched WGS datasets (e.g., MuTect2 vs. Strelka2) using appropriate correlation coefficients.

### Stratification by LoF

Patients were stratified according to LoF status (e.g., LoF = 1 versus Wild Type) to investigate the association between gene inactivation and microhomology-associated variant burden. Using WGS data (Strelka2), absolute counts of microhomology-mediated deletions were compared between groups using the Wilcoxon rank-sum test. Results were visualized using volcano plots, with log_2_ fold change between cohorts on the x-axis and statistical significance (-log_10_(adjusted p-value)) on the y-axis.

For selected genes showing strong associations, the effect of varying the parameter *M_lt_* on deletion classification and microhomology length distribution was further evaluated across LoF groups. Additionally, results for selected genes in both LoF and Wild Type groups were presented using boxplots.

### Survival and hazard ratio analysis

For survival analyses, patients were stratified into high- and low-burden groups by dividing them according to the median proportion of individual variant classes (DEL, INS, MNVs). To estimate hazard ratios for selected genomic variables, a series of separate Cox proportional hazards models were fitted. Specifically, each genomic feature was evaluated independently in its own model, with age and tumor stage consistently included as covariates to adjust for clinical confounding. Predictive performance for each model was evaluated using the concordance index (C-index). The analyzed variables included absolute counts and proportions of microhomology-mediated events, tandem repeat-associated events, insertions, and MNVs, as well as the average microhomology length.

Survival probabilities were estimated using Kaplan-Meier curves, and differences between groups were assessed using the log-rank test. Forest plots were generated to summarize HR estimates for features associated with deletions, insertions, and MNV variants.

### Pan-cancer and RMST analysis

The pan-cancer analysis assessed the relative contribution of different mechanisms generating deletions (classified as microhomology-associated, repeat-mediated, or other mechanisms) based on WGS data across multiple cancer types. The total number of deletions identified in each tumor type was quantified with stratification by deletion category (using a broken Y-axis for cohorts with exceptionally high mutation burdens), and the distribution of microhomology-associated deletion proportions across patients within each cohort was visualized using boxplots.

Cross-platform concordance was evaluated by assessing the correlation between cohort-level median levels of microhomology-mediated deletions estimated from WGS and WES data. Furthermore, the concordance of microhomology mediated deletion proportions was analyzed in matched TCGA WGS and WES samples at the individual patient level.

To comprehensively assess survival outcomes, restricted mean survival time (RMST) analysis was performed across all pan-cancer cohorts. Patients were stratified according to the median proportion of microhomology-associated deletions within each cancer type. Significant associations (p < 0.05) between microhomology-associated deletions and survival outcomes were summarized and highlighted using a cross-cohort volcano plot.

Using genomic data from the TCGA project, a Generalized Linear Model (GLM) analysis was conducted to identify the gene-level LoF alterations associated with the microhomology-mediated deletions. For this purpose, a quasi-binomial GLM was employed, with the response modeled as *cbind(microhomology-dependent deletion counts, non-microhomology-dependent deletion counts)*. Starting from an initial pool of 19,435 genes, a unified LoF status was established for each gene by combining SVs, SNVs, and CNVs into a single variable. To avoid bias from cohort-specific events and focus exclusively on alterations with pan-cancer relevance, the total number of mutated samples per gene was calculated, and the maximum proportion of these mutations attributed to any single cancer type was determined. Genes were retained for downstream analysis only if they were LoF in more than 100 samples overall, and provided that no single cancer type contributed to more than 50% of a given gene’s total LoF count. For the filtered data, a preliminary GLM based on the overall LoF status was constructed, adjusting for cancer type. This allowed for the selection of 2678 genes showing high statistical significance (adjusted p < 0.001) and a strong effect size (|log_10_(Odds Ratio)| > 0.2). To ensure statistical robustness and prevent bias driven by a single cancer type, genes were further filtered to retain only those with Loss of Function (LoF) mutations in more than 10 samples and where the dominant cancer type accounted for no more than 50% of the mutated samples. Finally, for the selected genes, a GLM was performed, again adjusting for cancer type. In this final model, the LoF variable was separated back into individual mutation types (SV, SNV, CNV) to estimate their associations with microhomology-associated deletion burden.

## Results

### Optimization of microhomology length (parameter *M_lt_*) for detection of MMEJ signatures in ovarian cancer

To ensure robust quantification of microhomology-associated events, we first optimized the threshold for the minimum length of microhomology between the deletion and the post-deletion sequence (parameter *M_lt_*). We found that estimates of the microhomology-associated deletion signal remained stable within the range of *M_lt_* = 2-4, indicating that this interval provides a robust classification window. However, increasing beyond this range introduced an overly stringent classification criterion, resulting in a rapid loss of detectable events and reduced statistical stability of downstream analyses.

An analysis designed to distinguish the true biological mechanism from random genomic background (signal-to-noise ratio) revealed a strong dependence on the microhomology threshold. For short matching requirements (low values of parameter *M_lt_*), the actual number of observed homologous sequences was substantially higher than expected by chance alone (Figure 2A). The ratio of observed to expected events (estimated empirically using a genomic background window shifted by +100 nt) reached its maximum at *M_lt_* = 2-3. In contrast, increasing *M_lt_* resulted in a sharp decline of the observed-to-expected ratio, falling below 1 and approaching zero. This indicates underrepresentation of detectable microhomology-associated events under overly stringent thresholds, ultimately leading to a loss of biologically informative signal (Figure 2A).

**Figure 2.**
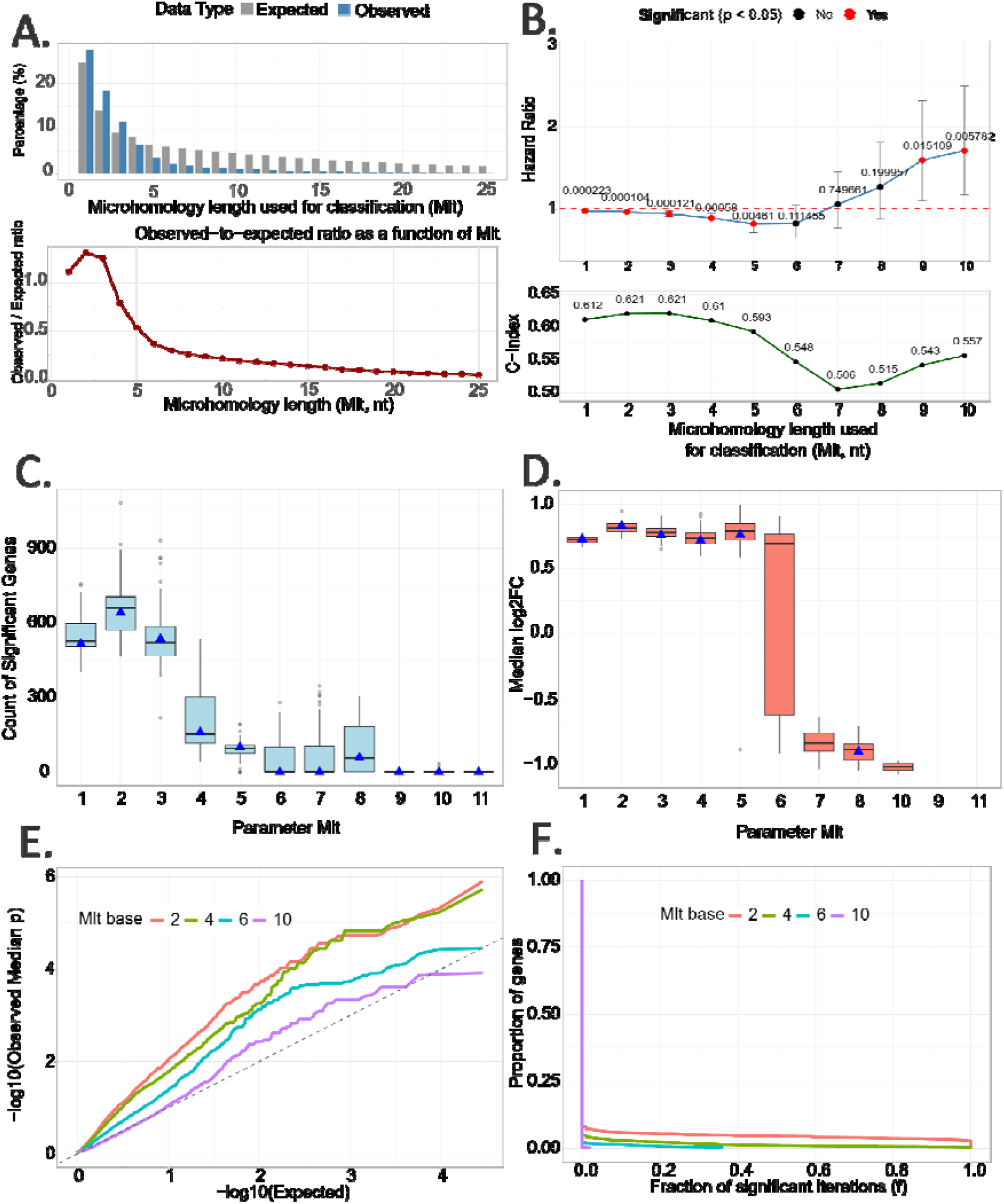
Selection of the Optimal Microhomology Length Threshold for Robust Detection of Microhomology-mediated Deletions. A. SNR analysis of microhomology-mediated deletion detection across increasing microhomology length thresholds (M_lt_). The upper panel shows the observed (blue) and expected (gray) cumulative percentages of MH events relative to all deletions, with expected values derived from a shifted genomic background window (+100 nt). The lower panel shows the observed-to-expected ratio as a function of M_lt_. B. Prognostic optimization of M_lt_ in ovarian cancer. The upper panel shows hazard ratios (HR) for the percentage of Microhomology-mediated deletions across M_lt_ values, with statistically significant associations highlighted in red (p < 0.05) Cox analysis. The emergence of statistical significance at specific values of M_lt_ suggests that these thresholds may more effectively filter sequencing noise, potentially capturing microhomology-mediated deletions. The lower panel shows C-index for survival models across parameter settings. C. Stability of gene-level LoF associations under downsampling. Boxplots represent the distribution of the number of significant genes across 100 bootstrap iterations; blue triangles indicate baseline (non-downsampled) results. D. Stability of effect size estimates (median log2 fold change) across M_lt_ values under downsampling. Blue triangles indicate baseline estimates. E. QQ-plot comparing observed and expected distributions of -log10(p) values after normalization for sample size across selected M_lt_ thresholds (M_lt_ = 2, 4, 6, 10). F. Cumulative distribution function (CDF) of gene-level reproducibility across downsampling iterations, showing the proportion of genes remaining significantly associated (p < 0.05) across bootstrap replicates for each M_lt_ value.

Next, we assessed the impact of parameter *M_lt_* on the performance of prognostic models in the studied cohort. Values within the range of *M_lt_* = 2-5 maximized the predictive performance of survival models, as reflected by the highest C-index values shown in the lower panel of Figure 2B. Increasing the threshold resulted in a monotonic decline in model performance. Although statistically significant hazard ratios were observed for selected higher values of *M_lt_*, these estimates were highly unstable due to the drastic reduction in the number of identified variants (Figure 2B).

The reliability of the selected threshold was further evaluated using a downsampling analysis. We found that for low and intermediate values of *M_lt_*, both the ability to identify Loss of Function (LoF) genes (Figure 2C) and the estimated magnitude of this effect (log_2_ fold-change, Figure 2D) remained highly stable. In contrast, requiring excessively long matching microhomology tracts through the application of higher *M_lt_* values led to a reduction in the number of detected genes and an artificial attenuation of effect sizes. These findings demonstrate that an overly stringent threshold does not improve accuracy but instead causes an irreversible loss of genuine biological signal.

These conclusions were further supported by a statistical assessment using QQ-plots (Figure 2E). Values of *M_lt_*= 2 and *M_lt_*= 4 showed the strongest deviation from the null expectation, suggesting that these settings may enable the algorithm to detect the greatest number of potential biological associations. Furthermore, bootstrap resampling analysis (Figure 2F) demonstrated that these thresholds were robust, with the algorithm consistently identifying similar sets of significant genes across resampling iterations. In contrast, higher values of *M_lt_* resulted in a marked decline in result reproducibility, confirming their statistical instability.

### Stability of Detection of Microhomology-mediated Deletions and the Impact of Sequencing Technology on MMEJ Signature Identification

After establishing *M_lt_* = 2 as the optimal threshold to maximize the sensitivity and stability of microhomology-mediated deletion detection, the robustness of this calibrated metric was evaluated across different technological environments. Given that short-variant detection is highly dependent on the statistical assumptions and quality filters implemented by variant-calling algorithms, assessing the consistency of this metric across sequencing platforms and analytical software was essential. Consequently, a comparative analysis of leading variant callers was performed using WES and WGS data.

Comparison of the results revealed substantial differences in the distribution of deletion classes reported by individual algorithms (Figure 3A). The relative proportions of Microhomology-mediated, repeat-mediated, and other deletion classes varied considerably depending on the variant caller used. This variability was particularly pronounced in WES data because algorithms struggle with the incomplete genomic context, whereas WGS analyses produced substantially more consistent and reproducible profiles by offering a full sequence context, indicating greater reliability for the identification of microhomology-mediated deletion signatures.

**Figure 3.**
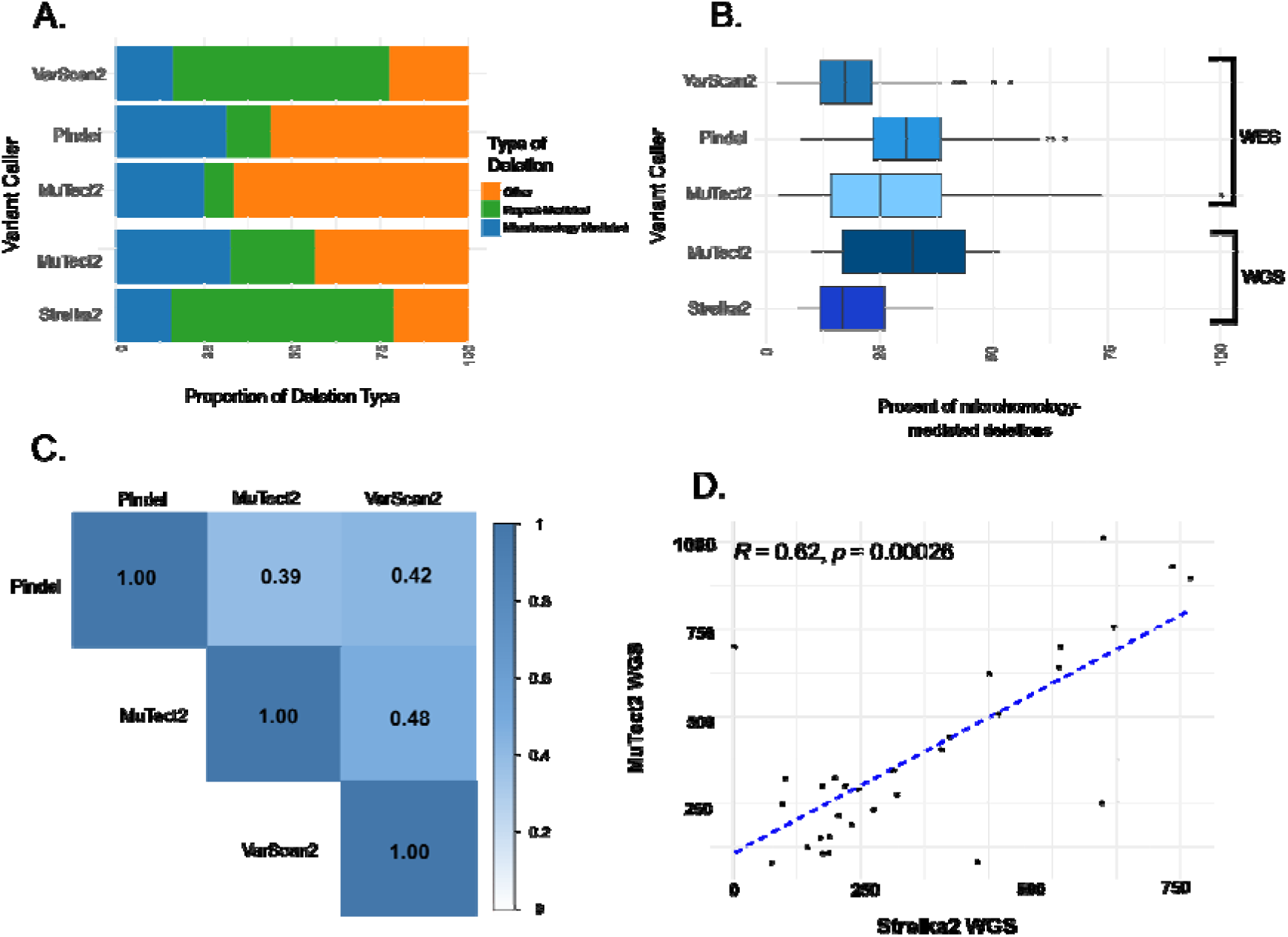
Evaluation of microhomology-mediated deletion detection and compatibility between variant-calling algorithms in WES and WGS datasets. A. Stacked bar plot showing the relative proportions of deletion classes identified by different variant-calling algorithms, including VarScan2 WES, Pindel WES, MuTect2 WES, MuTect2 WGS, and Strelka2 WGS. Deletions were classified as microhomology-mediated, repeat-mediated, or other events. B. Boxplots illustrating the distribution of the percentage of microhomology-mediated deletions detected by each variant caller across analyzed samples. The plots highlight differences in the estimated contribution of MMEJ-associated events between WES- and WGS-based analytical pipelines. C. Correlation matrix comparing the concordance of microhomology-mediated deletion proportions estimated by three variant callers applied to WES data (Pindel, MuTect2, and VarScan2). Pairwise correlation coefficients ranged from 0.39 to 0.48, indicating limited agreement among algorithms. D. Scatter plot comparing the number of MMEJ-associated deletions identified independently by MuTect2 and Strelka2 in WGS datasets. A moderate positive correlation was observed (R = 0.62, p = 0.00026), supporting improved reproducibility of MMEJ detection in whole-genome sequencing data.

The distribution of microhomology-mediated deletion frequencies across individual samples further confirmed this variability (Figure 3B). Although all tools detected a measurable proportion of deletions consistent with microhomology-mediated repair, the estimated frequencies differed substantially between algorithms. These discrepancies likely reflect differences in underlying detection models and quality-filtering strategies, which directly affect short indels and the accurate classification of deletion events.

To quantitatively assess the consistency of WES-based approaches, correlation analyses were performed between the proportions of microhomology-mediated deletions identified by different variant callers for each patient (Figure 3C). Agreement among the evaluated tools (Pindel, MuTect2, and VarScan2) was limited, with correlation coefficients ranging from R = 0.39 to R = 0.48. These findings indicate that the estimation of microhomology-mediated deletion burden from exome sequencing data is strongly influenced by the choice of variant-calling algorithm, posing a significant challenge for the standardization and reproducibility of WES-based analyses.

Analysis of WGS data using MuTect2 and Strelka2 demonstrated higher concordance between algorithms (Figure 3D). The numbers of microhomology-mediated deletions independently identified by both tools showed a markedly stronger and highly statistically significant correlation (R = 0.62, p = 0.00026). This observed reproducibility implies that comprehensive genome-wide coverage could offer a more reliable baseline for studying DNA repair-associated mutational patterns. Given the limited consistency observed in exome-based analyses, WGS emerges as the preferred and substantially more precise platform for mechanistic studies of deletion formation and for the identification of deletion signatures consistent with microhomology-mediated repair. Furthermore, an assessment of overall variant frequencies revealed that deletions are the most abundant variant type, and there is a positive, though not strong, correlation between the total number of deletions and MNVs (Supplementary Figure 1).

### Gene-level determinants of microhomology-associated deletion burden in ovarian cancer

Following the optimization of the microhomology length threshold (parameter *M_lt_* = 2, corresponding to sequence concordance within the deletion region and immediately downstream of it) and the confirmation of detection stability using WGS data, we proceeded to verify the tool’s performance in a clinical context by analyzing an ovarian cancer cohort. Specifically, we investigated whether the LoF in key genes directly correlates with an increased burden of microhomology-mediated deletions, thereby serving as a reliable indicator of underlying DNA repair deficiencies.

The analysis revealed that ovarian cancer patients showing LoF in specific tumor suppressor genes presented a highly significant overrepresentation of microhomology-dependent deletions (Figure 4A). As detailed in Supplementary Table 1, inactivating mutations in genes such as RB1, CCDC122, BRI3BP, DHX37, and LACC1 were associated with positive log_2_ fold change values, indicating a higher number of microhomology-mediated deletions in patients with LoF compared to wild type alleles. This finding suggests that LoF status in these genes is associated with increased microhomology-associated deletion burden.

**Figure 4.**
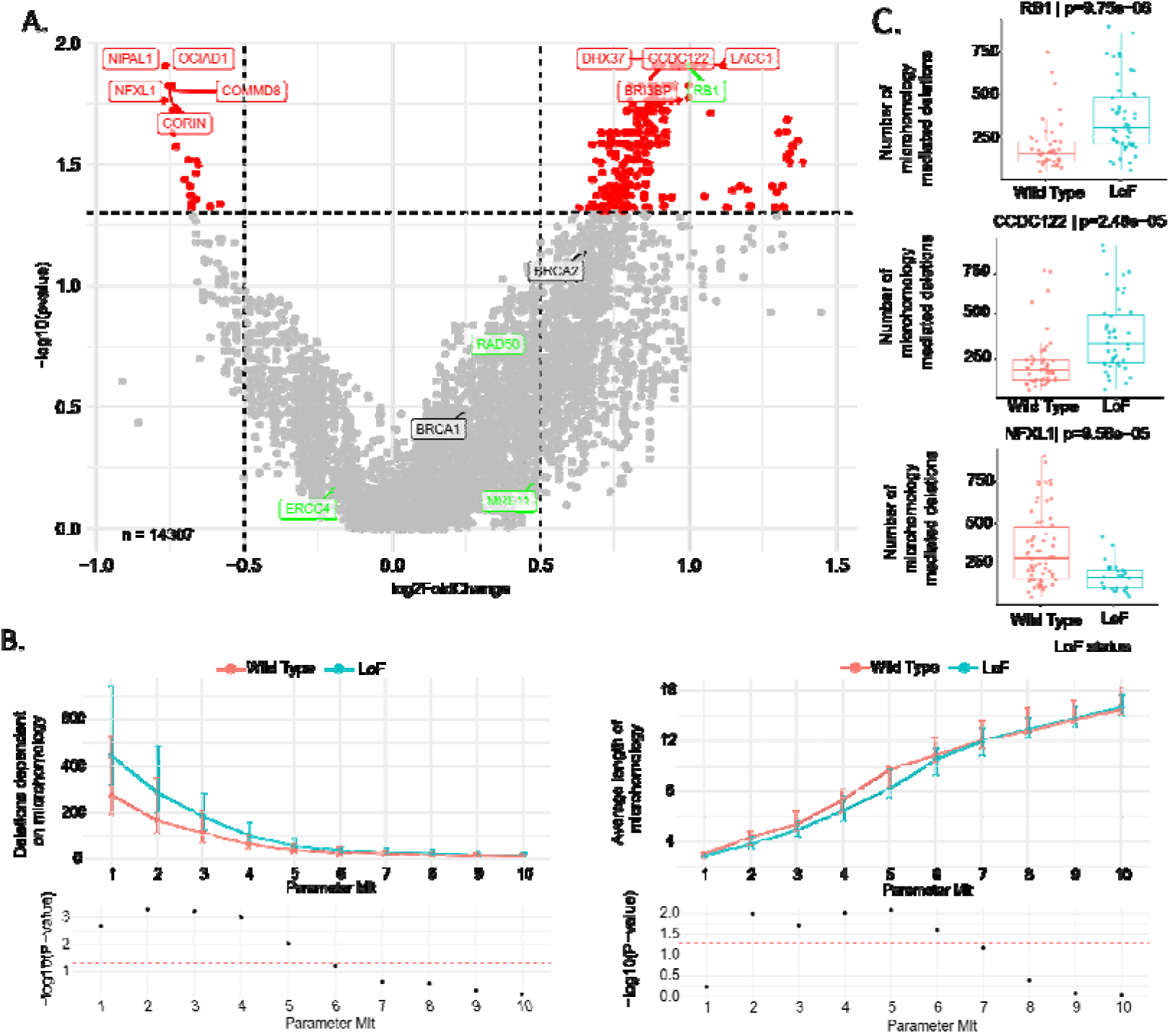
Gene-Level Associations Between Loss of Function Events and Microhomology-Mediated Deletions in Ovarian Cancer (OV). A. Volcano plot identifying genes enriched for microhomology-mediated deletions. The x-axis represents the log_2_ fold change between LoF and Wild Type groups, while the y-axis shows statistical significance -log_10_(p-value) derived from Wilcoxon tests. Significantly enriched genes are highlighted in red. B. Optimization of the MMEJ detection algorithm using the parameter M_lt_ (minimum number of matching nucleotides downstream of the deletion), evaluated for the RB1 gene. The top panel shows changes in the number of detected deletions with increasing M_lt_, stratified by LoF status. The middle panel illustrates changes in average microhomology length, while the bottom panels display corresponding p-values across parameter settings. C. Boxplots comparing the absolute number of microhomology-mediated deletions between LoF and Wild Type groups for selected significantly associated genes.

A pivotal example confirming the method’s biological sensitivity is RB1, a gene encoding a tumor suppressor protein responsible for cell cycle control. The applied classifier, utilizing the M_lt_ = 2 threshold, precisely identified an elevated microhomology-associated signature burden in patients with the inactivation of this gene (Figures 4B and 4C) (36). The presence of this gene in the high-burden group confirms that the optimized measure captures a sequence-based signature consistent with increased use of alternative repair pathways, which are upregulated in response to the failure of primary mechanisms such as HR (37).

Simultaneously, the application of the highly sensitive *M_lt_* = 2 threshold demonstrated that individual genes can exert varied effects on the mutational profile. Alongside the aforementioned genes, we identified a distinct set of genes (including NIPAL1, NFXL1, CORIN, OCIAD1, and COMMD8 in Supplementary Table 1) for which preserved function (Wild Type) was associated with a significantly higher level of microhomology-mediated deletions, as reflected by negative log_2_ fold change values.

However, our primary mechanistic focus remains on cases where gene inactivation actively drives genomic instability. This critical dynamic is prominently illustrated by the analysis of genes such as RB1 and CCDC122 (Figure 4C), where patients with impaired gene function (the LoF group) exhibited a strikingly higher microhomology-associated deletion burden compared to those with preserved functional activity.

### Prognostic Significance of Microhomology-Associated Indel and MNV Signatures in Ovarian Cancer

To evaluate the clinical relevance of the identified mutational signatures, we investigated the relationship between ovarian cancer patient outcomes and genomic burden across different classes of variants (deletions, insertions, and MNVs), adjusting for patient age and cancer stage, with particular emphasis on their association with microhomology-mediated mechanisms.

Stratification of patients based on the median proportion of microhomology-dependent deletions revealed a strong and highly statistically significant association with overall survival (Figure 5A). Patients characterized by a high burden of microhomology-mediated deletions exhibited significantly longer survival compared to the low-burden group (log-rank test, p < 0.0001). This indicates that deletion-based MMEJ signatures represent a prognostic factor in this cancer type.

**Figure 5.**
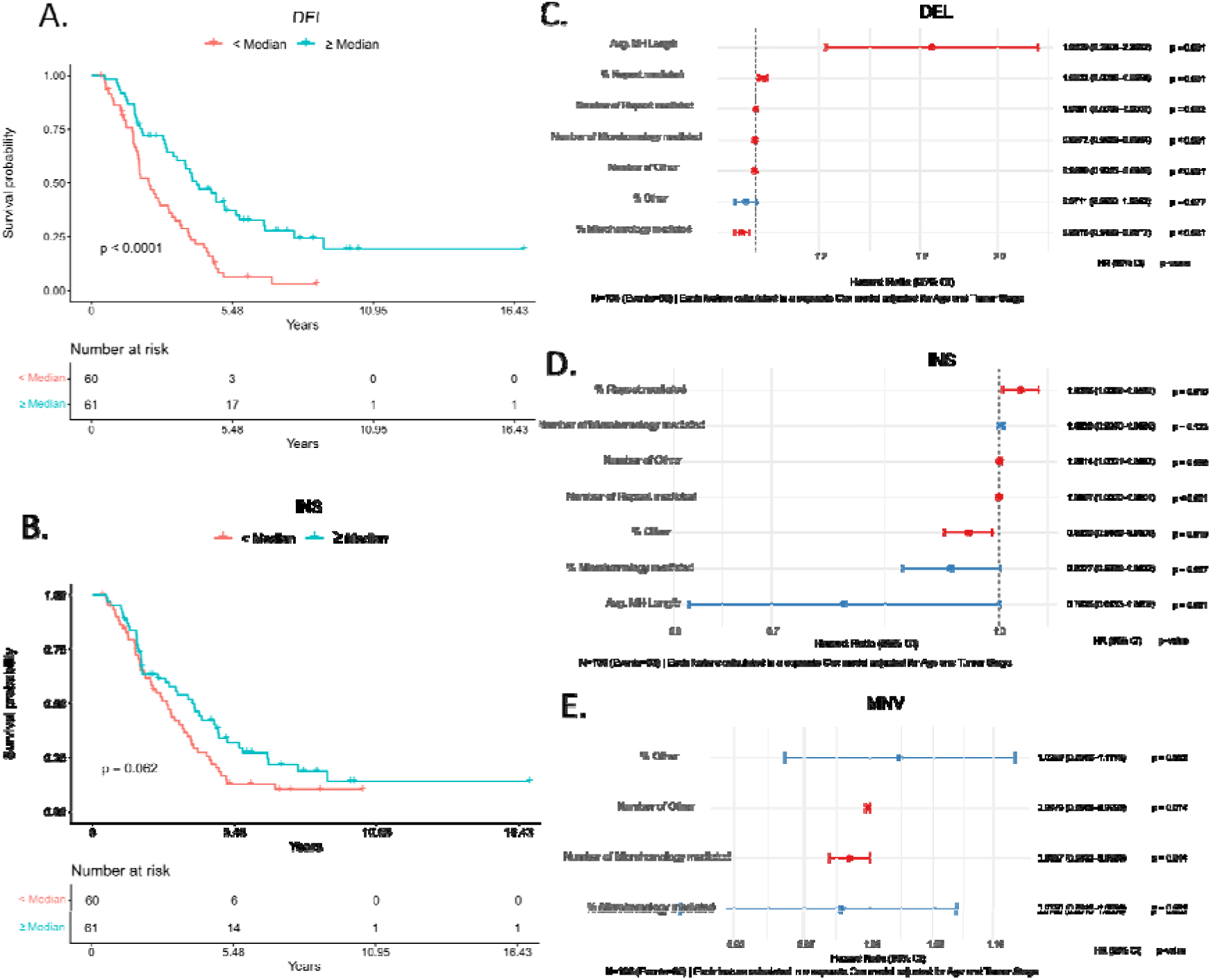
Prognostic impact of microhomology-associated indel in ovarian cancer. A. Kaplan-Meier survival curves stratifying ovarian cancer patients into high- and low-burden groups based on the median proportion of microhomology-mediated deletions. B. Kaplan-Meier survival analysis stratified by the median proportion of microhomology-mediated insertions. C. Forest plot of hazard ratios (HR) for deletion-associated features, including average microhomology length, proportion of Microhomology-mediated deletions, and absolute deletion counts. D. Forest plot of hazard ratios for insertion-associated features, highlighting significant prognostic variables. E. Forest plot summarizing hazard ratios for MNV-associated features, including MH-related burden and class-specific variant counts.

In contrast to deletions, an analogous stratification based on the median proportion of microhomology-dependent insertions showed only weak separation of survival curves, which did not reach statistical significance (Figure 5B, p = 0.062). Additionally, we evaluated the broader prognostic impact of general indel burden using WES data; similar survival analyses were performed based on the overall percentage of deletions and insertions in the genome (Supplementary Figure 2) as well as their cumulative length (Supplementary Figure 3).

To further investigate the contribution of individual features to mortality risk, each parameter was evaluated in separate Cox regression models adjusted for clinical covariates (Age and Tumor Stage).

Analysis of deletion-related features (Figure 5C) revealed that average microhomology length (Avg. MH Length) was the only parameter significantly associated with an increased mortality risk. Conversely, the remaining statistically significant metrics, including the overall proportion of both microhomology- and repeat-mediated deletions, as well as the absolute counts of all deletion classes, were associated with improved survival (HR < 1). The proportion of unclassified deletions was the only feature that did not show a statistically significant effect.

Greater heterogeneity was observed in the adjusted analysis of insertion-related features (Figure 5D). Notably, repeat-mediated insertion burden (both percentage and count) was significantly associated with worse clinical outcomes. In contrast to deletions, microhomology-driven insertion features did not reach statistical significance. Furthermore, while the absolute count of unclassified insertions showed a significant increase in risk, a higher overall proportion of these events was associated with decreased mortality hazard.

Finally, analysis of MNV-based features (Figure 5E) demonstrated that higher absolute counts of both microhomology-mediated and unclassified MNV events were significantly associated with improved survival outcomes. However, the continuous proportional metrics for these classes did not show stable, statistically significant associations with patient survival.

### Pan-cancer Landscape of Microhomology-associated Deletion Burden

After characterizing microhomology-associated deletion patterns in ovarian cancer, we next extended the analysis to a broader TCGA pan-cancer cohort. This analysis revealed substantial variability in the relative contribution of microhomology-associated deletions across cancer types.

Tumors such as testicular germ cell tumors (TGCT), pheochromocytoma and paraganglioma (PCPG), and uveal melanoma (UVM) exhibited the highest relative proportion of microhomology-dependent deletions, often exceeding 25% of all analyzed events (Figure 6A and 6C). In contrast, cohorts such as endometrial carcinoma (UCEC) were dominated by repeat-mediated and other alternative deletion mechanisms. These findings suggest that microhomology-associated deletion burden is strongly shaped by tumor-specific genomic context.

**Figure 6.**
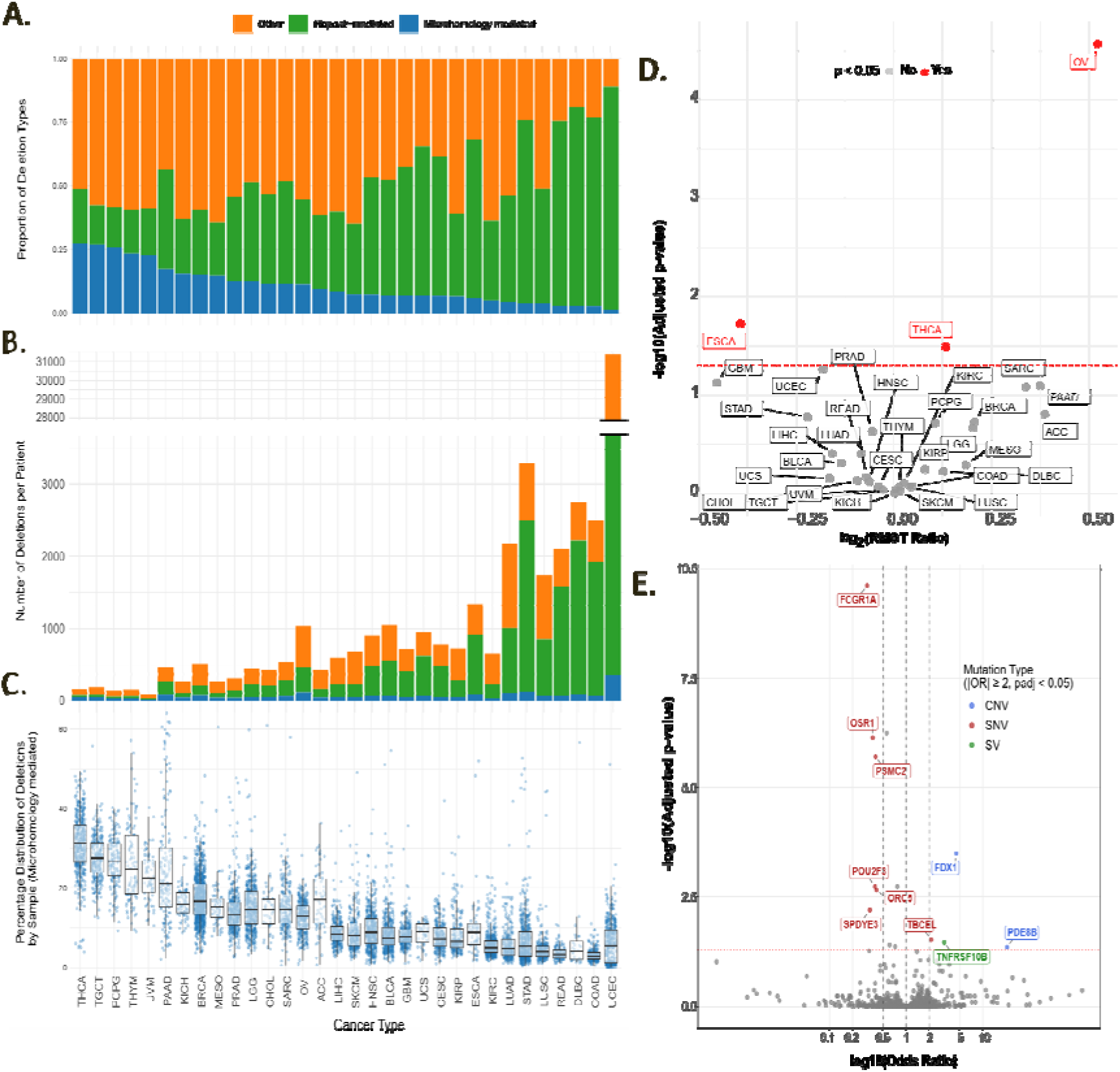
Landscape and clinical relevance of microhomology-mediated deletions across human cancers. A. Stacked bar plot showing the relative contribution of different deletion-generating mechanisms in WGS datasets across cancer types. Deletions were classified as microhomology-mediated (blue), repeat-mediated (green), or other mechanisms (orange). B. Total number of deletions identified in each cancer type, stratified according to the same mechanistic categories as in panel A. A broken y-axis was applied to accommodate highly mutated outlier cohorts. C. Boxplots illustrating the distribution of the proportion of microhomology-mediated deletions across individual patient samples within each cancer cohort. D. Volcano plot summarizing Restricted Mean Survival Time (RMST) analysis across PanCancer cohorts. Patients were stratified according to the median proportion of microhomology-mediated deletions within each cancer type. Significant associations between microhomology-associated deletion burden and survival outcome are highlighted (p < 0.05), including esophageal carcinoma (ESCA), thyroid carcinoma (THCA), and ovarian cancer (OV). E. Volcano plot showing the impact of Loss of Function (LoF) variants on the proportion of microhomology-mediated deletions, adjusted for cancer type. Colors highlight specific mutation types (SNV, CNV, SV) with a strong and statistically significant effect (adjusted p < 0.05 and OR ≥ 2 or ≤ 0.5).

Importantly, it was demonstrated that the relative contribution of the MMEJ signature does not directly reflect the total deletion burden (Figure 6B). Highly unstable tumor types, such as endometrial carcinoma (UCEC) or stomach adenocarcinoma (STAD), accumulated extremely large numbers of variants while maintaining a relatively low proportion of microhomology-dependent events. This observation suggests that extensive genomic instability in these cancers is primarily driven by other mutational processes or genomic instability mechanisms not captured by the microhomology-associated deletion signature.

As part of methodological validation, the utility of both sequencing platforms was compared using TCGA data. It was shown that WES allows for a relatively accurate estimation of total deletion burden at the level of aggregated cohorts, as supported by a strong correlation with WGS results (R = 0.89, Supplementary Figure 4). However, moderate concordance in paired samples at the individual patient level (Supplementary Figure 4, R = 0.59) indicates that the limited genomic coverage of WES does not allow precise profiling of DNA repair signatures in individual cases. These results support WGS as the preferred platform for the analyses of microhomology-associated deletion signatures across patient samples.

Multi-cohort survival analysis (Restricted Mean Survival Time, RMST) demonstrated that the prognostic relevance of Microhomology-mediated deletions is highly tissue-context dependent (Figure 6D). The impact of elevated microhomology-associated deletion levels on patient survival was significant only in selected tumor types. In addition to OV we observed significant associations in esophageal carcinoma (ESCA) and thyroid carcinoma (THCA). These results indicate that although MMEJ-related deletion signatures have strong potential as clinically informative biomarkers, their prognostic value is not universal but instead remains strictly specific to particular tumor types.

Although the prognostic value of microhomology-mediated deletions varied across tumor types, we next investigated whether their burden was linked to recurrent gene-level Loss of Function events shared across all analyzed cancers. To this end, we applied a Generalized Linear Model (GLM) adjusted for cancer type, which allowed us to assess the impact of LoF mutations in individual genes on the proportion of these deletions, independent of baseline differences between tumors. The volcano plot (Figure 6E) highlights variants (SNV, CNV, SV) with a highly significant and strong effect (adjusted p-value < 0.05 and Odds Ratio ≥ 2 or ≤ 0.5). Detailed statistics and functional annotations for these significant variants are provided in Supplementary Table 2. Genes with the highest odds ratios (e.g., FDX1 and PDE8B) represent promising candidates for universal pan-cancer markers, whose Loss of Function was associated with increased burden of deletions consistent with MMEJ-mediated repair.

## Discussion

Identification of mutations potentially generated through MMEJ pathway constitutes an important component of molecular tumor profiling. Our analyses demonstrated that WES lacks sufficient resolution for the accurate assessment of MMEJ-associated mutational signatures. Reliable detection of microhomology-associated deletion enrichment is better achieved using WGS, where the total number of detectable deletions is much higher and is not restricted to coding or capture-targeted regions, which may be subject to stronger selective constraints. A critical parameter determining the reliability of the obtained results is the appropriate calibration of the minimum microhomology length threshold between the deletion sequence and its surrounding genomic context. Setting this threshold at *M_lt_* = 2 prevents the loss of biologically relevant mutations that occur when overly stringent filtering criteria are applied.

LoF alterations in DNA repair-related genes can induce shifts in DNA repair pathway utilization in cancer cells. Our findings corroborate previous reports showing that RB1 loss is associated with impaired ssDNA resection and increased reliance on the MMEJ pathway at the expense of HR (38). In OV, LoF alterations in genes such as CCDC122, LACC1, BRI3BP, and DHX37 are associated with a similar increase in cellular dependence on microhomology-mediated DNA repair.

In prostate adenocarcinoma (PRAD), dysfunction of CCDC122 and LACC1 co-occurs with MYC oncogene hyperactivation, contributing to elevated replication stress. Breakpoint analysis at the MYC locus indicates that local rearrangements are consistent with Microhomology-mediated repair mechanisms (39). This results in increased cellular dependence on this pathway for the repair of double-strand breaks, providing a rationale for investigating Polθ and PARP inhibitors as potential therapeutic strategies (40).

Similar pathogenetic relationships are observed in HCC. Integration of HBV viral DNA into the host genome, an essential event in hepatocarcinogenesis, is in these tumors tightly dependent on MMEJ pathway activity under conditions of replication stress (41). The clinical relevance of these processes is supported by a negative correlation between patient survival and elevated DHX37 expression (42), as well as an association between BRI3BP overexpression and higher tumor stage (43). The occurrence of these alterations creates an intracellular environment that promotes tumor progression, strongly dependent on microhomology-based repair mechanisms (44).

Pan-cancer analyses using TCGA data indicate a tissue-specific pattern of tumor dependence on the MMEJ pathway. For example, TGCT comprise subtypes with distinct developmental potential that exhibit differential requirements for double-strand break repair pathways, including MMEJ (45). Moreover, PCPG are characterized by an intrinsically high fraction of MMEJ-associated deletions, in contrast to UCEC, which displays alternative mutational profiles.

Quantification of deletions consistent with MMEJ-mediated repair also shows prognostic relevance. In OV, the accumulation of microhomology-associated deletion burden was associated with improved overall survival. Prognostic associations of the MMEJ pathway have also been reported in ESCA and THCA. In THCA, elevated Polθ expression correlates with disease progression dynamics, PARP inhibitor resistance, and poor prognosis (46).

A plausible biological explanation for the association between higher microhomology-associated deletion burden and improved survival in OV is that this signal may reflect an underlying HRD-like repair context rather than a direct protective effect of MMEJ itself. Ovarian tumors with impaired high-fidelity double-strand break repair, particularly HR deficiency, may increasingly rely on error-prone backup pathways such as MMEJ (47). Consequently, elevated microhomology-associated deletion burden may serve as a molecular footprint of defective HR activity (48). At the same time, this repair-deficient state may increase tumor sensitivity to DNA-damaging therapies, particularly platinum-based chemotherapy and PARP inhibition, thereby providing a potential explanation for the improved survival observed in patients with higher microhomology-associated deletion burden. However, this interpretation remains indirect and requires validation in cohorts with detailed treatment annotations and established HRD biomarkers.

When investigating differences between cancer types, our GLM analysis identified several important genes, including FDX1 and PDE8B, whose Loss of Function correlates with increased microhomology-associated deletion burden. These associations may be biologically plausible, although indirect. FDX1 is primarily a regulator of cuproptosis and mitochondrial metabolism (49), it also influences the Hippo signaling pathway. Dysregulation of this pathway results in the activation and nuclear translocation of the YAP/TAZ transcriptional coactivators, thereby promoting cell survival (50). This is particularly relevant in the context of DNA damage, as YAP/TAZ drive the transcription of genes involved in high-fidelity DNA repair pathways, including HR, nucleotide excision repair (NER), and base excision repair (BER), thereby preserving genomic stability (50,51). Consequently, loss of FDX1 function and the resulting disruption of Hippo signaling may impair HR activity. Under these conditions, cancer cells become increasingly reliant on microhomology-mediated end joining (MMEJ) to repair DNA double-strand breaks.

A similar indirect effect on genomic stability was observed for PDE8B, which was identified in our GLM analysis. PDE8B encodes a phosphodiesterase, and its Loss of Function impairs cAMP degradation, leading to its pathological accumulation within cells (52,53). Elevated intracellular cAMP strongly activates the protein kinase A signaling axis (cAMP-PKA-pCREB), which enhances poly(ADP-ribose) polymerase (PARP) activity in response to DNA double-strand breaks (54). Because MMEJ is highly dependent on PARP1, hyperactivation of the cAMP-PKA-PARP axis resulting from PDE8B deficiency is expected to shift DNA repair toward this pathway. This provides a mechanistic explanation for why the model-predicted loss of PDE8B function is strongly associated with the accumulation of MMEJ-characteristic deletions. These suggest that the identified genes may serve as universal biomarkers associated with MMEJ-mediated deletions, independent of cancer type.

Overall, our findings support microhomology-associated deletion burden as a potentially informative marker of DNA repair deficiency and tumor-type-specific clinical behavior. However, the mechanistic interpretation of these signatures and their utility for patient stratification require validation in independent cohorts with harmonized WGS processing, treatment annotations, and orthogonal HRD measures.

## Funding

This research was funded by the Polish National Science Center grant 2021/41/B/NZ2/04134. MK work was supported in part by the Department of Defense through the Peer Reviewed Cancer Research Program under Award No. CA250098. Opinions, interpretations, conclusions, and recommendations are those of the author and are not necessarily endorsed by the Department of Defense.

## Acknowledgments

Calculations were carried out using the infrastructure of the Ziemowit computer cluster (www.ziemowit.hpc.polsl.pl) in the Laboratory of Bioinformatics and Computational Biology, The Biotechnology, Bioengineering and Bioinformatics Centre Silesian BIO-FARMA, created in the POIG.02.01.00-00-166/08 and expanded in the POIG.02.03.01-00-040/13 projects.

The results published here are in part based upon data generated by the TCGA Research Network (https://www.cancer.gov/tcga) and the International Cancer Genome Consortium (https://dcc.icgc.org/).

## Code Availability Statement

The code for MHDetect is available at https://github.com/darkoss566/MHDetect/.

## Data Availability

No new sequencing data were generated. All analyzed data were obtained from publicly available TCGA and ICGC resources, as described in the Data section.

## Conflict of Interest

The authors declare no conflicts of interest.

## Ethics statement

This study was based exclusively on previously generated, de-identified genomic and clinical data from The Cancer Genome Atlas (TCGA) and the International Cancer Genome Consortium (ICGC). No new human participants were recruited, no new biospecimens were collected, and no new sequencing data were generated. The original TCGA and ICGC studies obtained ethical approval and informed consent from participants according to their respective institutional and consortium-level procedures. Therefore, additional institutional ethics approval was not required for this secondary analysis. All data were accessed and analyzed in accordance with TCGA, GDC, and ICGC data-use policies. Where controlled-access data were used, access was obtained under the applicable data-use agreements and consortium access policies.

## Glossary

ACC: Adrenocortical carcinoma - A rare cancer of the adrenal cortex
alt-NHEJ: Alternative nonhomologous end joining
BLCA: Bladder urothelial carcinoma - The most common type of bladder cancer
BRCA: Breast invasive carcinoma - A broad term for invasive breast cancers
BRCA1/2: Breast cancer type 1/2 susceptibility proteins
C-NHEJ: Canonical nonhomologous end joining
CESC: Cervical squamous cell carcinoma and endocervical adenocarcinoma - Cancers of the cervix
CHOL: Cholangiocarcinoma - Cancer of the bile ducts
COAD: Colon adenocarcinoma - The most common form of colon cancer
CtIP: C-terminal binding protein interacting protein
DLBC: Diffuse large B-cell lymphoma - A common type of non-Hodgkin lymphoma
DSBs: DNA double-strand breaks
ERCC1/XPF: Excision repair cross-complementation group 1/Xeroderma pigmentosum complementation group F
ESCA: Esophageal carcinoma - Cancer of the esophagus, often of squamous cell or adenocarcinoma type
FDA: Food and Drug Administration
FEN1: Flap structure-specific endonuclease 1
GBM: Glioblastoma multiforme - A highly aggressive type of brain cancer
HMCES: Hydroxymethylcytosine epithelial splicing protein
HNSC: Head and neck squamous cell carcinoma - Cancer of the head and neck, often associated with smoking or HPV
HR: Homologous recombination
HRD: Homologous recombination deficiency
KICH: Kidney chromophobe carcinoma - A rare type of kidney cancer
KIRC: Kidney renal clear cell carcinoma - The most common type of kidney cancer
KIRP: Kidney renal papillary cell carcinoma - A type of kidney cancer with papillary architecture
Ku70/80: Ku70 and Ku80 proteins (involved in DNA repair)
LAML: Acute myeloid leukemia - A cancer of the blood and bone marrow
LGG: Lower grade glioma - Less aggressive brain tumors compared to glioblastoma
LIHC: Liver hepatocellular carcinoma - The most common primary liver cancer
LIG3/XRCC1: DNA ligase III/X-ray repair cross-complementing protein 1
LUAD: Lung adenocarcinoma - A common subtype of non-small cell lung cancer
LUSC: Lung squamous cell carcinoma - Another major subtype of non-small cell lung cancer
MESO: Mesothelioma - A cancer of the mesothelium, often related to asbestos exposure
MMEJ: Microhomology-mediated end joining
MRN: Mre11-Rad50-Nbs1 complex
MNVs: Multinucleotide variants
NHEJ: Nonhomologous end joining
OV: Ovarian serous cystadenocarcinoma - The most common type of ovarian cancer
PAAD: Pancreatic adenocarcinoma - A common type of pancreatic cancer
PARP-1: Poly(ADP-ribose) polymerase 1
PARPi: Poly(ADP-ribose) polymerase inhibitors
PCPG: Pheochromocytoma and paraganglioma - Tumors that arise from adrenal gland cells or nerve tissue
Polθ: DNA polymerase theta
PRAD: Prostate adenocarcinoma - The most common prostate cancer
RB: Retinoblastoma protein
READ: Rectum adenocarcinoma - The most common form of rectal cancer
RPA: Replication protein A
SARC: Sarcoma - A group of cancers that arise in bones and soft tissues
SKCM: Skin cutaneous melanoma - A type of skin cancer originating from melanocytes
ssDNA: Single-stranded DNA
STAD: Stomach adenocarcinoma - The most common type of stomach cancer
TGCT: Testicular germ cell tumors - Tumors originating from germ cells in the testicles
THCA: Thyroid carcinoma - Cancer originating in the thyroid gland, typically papillary or follicular
THYM: Thymoma - A rare tumor of the thymus gland
UCS: Uterine carcinosarcoma - A rare and aggressive form of uterine cancer
UCEC: Uterine corpus endometrial carcinoma - A common form of uterine cancer
UVM: Uveal melanoma - A rare type of melanoma occurring in the eye

## Notes

https://github.com/darkoss566/MHDetect/tree/master

https://gdc.cancer.gov/about-data/publications/pancanatlas

### Competing Interest Statement

The authors have declared no competing interest.

