## Supplementary File for "Microhomology-Driven Genomic Alterations in Cancer Genomes: Patterns, Prevalence, and Clinical Implications"

*Supplementary Figure 1 Summary of the frequency of occurrence of individual types of variants and the correlation between the number of deletions and multinucleotide changes.* ______ *2*
*Supplementary Figure 2 Kaplan-Meier survival analysis according to the percentage of deletions and insertions identified by the Mutect2, Pindel, and VarScan2 algorithms.* ______ *5*
*Supplementary Figure 3 Kaplan-Meier survival analysis according to the cumulative length of deletions and insertions identified by the Mutect2, Pindel, and VarScan2 algorithms.* ______ *6*
*Supplementary Figure 4 Landscape and clinical relevance of MMEJ-dependent deletions across human cancers.* *7*

*Supplementary Table 1 Genes exhibiting significant associations between LoF status and MMEJ-dependent mutation burden.* ______ *2*
*Supplementary Table 2 Genes identified by GLM analysis showing significant associations between specific mutation types and MMEJ burden.* ______ *8*

#### Mutational Landscape and Correlations Between Variant Types

The mutational landscape of the analyzed patient cohort is presented in Supplementary Figure 1. The bar plot (Panel A) illustrates the mean number of mutations per sample, stratified into deletions (DEL), insertions (INS), and multinucleotide variants (MNVs), with events further categorized according to their putative mechanisms of origin. The scatter plot (Panel B) demonstrates a positive linear correlation between the total number of deletions and the number of MNVs identified in samples from the same patients.

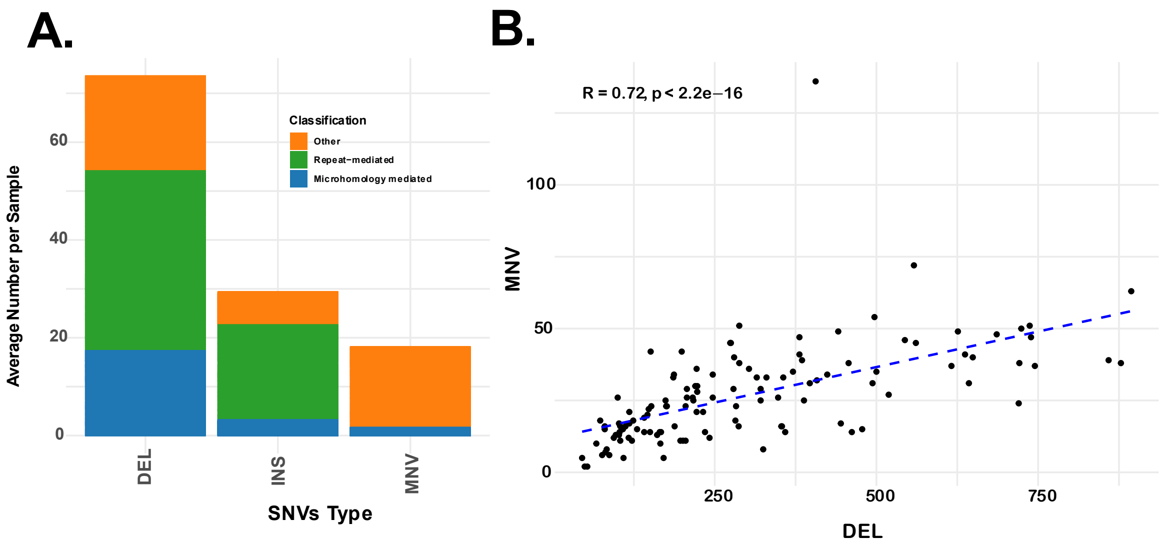


*Supplementary Figure 1 Summary of the frequency of occurrence of individual types of variants and the correlation between the number of deletions and multinucleotide changes. A. Bar chart showing the average number of mutations per sample, broken down by variant type: deletions, insertions, and multinucleotide variations. The individual components of the bars represent the classification of events based on their potential mechanism of occurrence. B. Scatterplot showing the relationship between the total number of deletions (X-axis) and the number of multinucleotide variations (Y-axis) identified within the same patient samples.*

#### Association Between LoF Status and MMEJ Mutations in Ovarian Cancer (OV)

Supplementary Table 1 summarizes the characteristics of genes whose Loss of function (LoF) status is significantly associated with the level of Microhomology-mediated deletions in patients with ovarian cancer (OV cohort). Positive $\log_2$ fold change values (e.g., for RB1, LACC1, and CCDC122) indicate an increased level of Microhomology-mediated deletions in patients harboring LoF alterations compared with the reference group (wild type). Conversely, negative values (e.g., for NIPAL1, NFXL1, and CORIN) indicate the opposite relationship, with higher levels of these deletions observed in patients retaining normal gene function. Statistical significance was assessed after correction for multiple testing using the Benjamini-Hochberg procedure (adjusted p-values). The table also includes the number of samples in each group (n) and a concise summary of the biological function of each gene.

*Supplementary Table 1 Genes exhibiting significant associations between LoF status and MMEJ-dependent mutation burden. The table presents the characteristics of genes for which Loss of function (LoF) significantly alters the level of MMEJ-dependent deletions. Positive $log_2$ fold change values indicate an increase in the analyzed variable within the LoF group compared to the reference (Wild Type) group. Conversely, negative $log_2$ fold change values reflect an inverse association, indicating a decrease in the mutation burden within the LoF group (i.e., preserved function is associated with a higher level of these deletions).*

*Abbreviations: BH - Benjamini-Hochberg procedure for p-value adjustment; LoF - Loss of Function; n - number of samples per group.*

| *Gene symbol* | *Adjusted* p-value *(BH)* | *$log_2$fold change* | *Group with LoF* (n) | *Group without LoF* (n) | *Function* | *LoF status* |
|---|---|---|---|---|---|---|
| *RB1* | 0.0124 | 1.00 | 49 | 42 | The gene encoding RB1 protein, tumor suppressor, controls the cell cycle, inhibits cell proliferation (1). | LoF |
| *CCDC122* | 0.0124 | 0.96 | 43 | 47 | Coiled-Coil Domain Containing 122 is overexpressed in prostate cancer compared to healthy tissue. Its differential expression and mutations are implicated in disease progression, while single nucleotide polymorphisms in this gene are associated with susceptibility to leprosy (2). | LoF |
| *BRI3BP* | 0.0124 | 0.91 | 32 | 58 | An RNA-binding protein upregulated in multiple cancers, including hepatocellular carcinoma, where it correlates with advanced tumor stage and poorer survival. It is functionally linked to cell cycle regulation, Rho GTPase activity, and copper homeostasis (3). | LoF |
| *DHX37* | 0.0124 | 0.91 | 32 | 58 | An RNA helicase involved in modifying RNA secondary structures, translation, splicing, and ribosome assembly. It acts as a functional regulator of CD8+ T cells, influencing antitumor immune responses. Its high expression is associated with worse prognosis in certain malignancies (4). | LoF |
| *LACC1* | 0.0124 | 1.02 | 42 | 48 | Encodes an oxidoreductase that promotes fatty-acid oxidation, inflammasome activation, reactive oxygen species (ROS) production, and anti-bacterial responses in macrophages. It exhibits altered expression in prostate cancer and, similarly to CCDC122, is linked to leprosy susceptibility (2). | LoF |
| *NIPAL1* | 0.0124 | -0.76 | 29 | 61 | Magnesium transporter, indirectly regulates uric acid metabolism; may participate in the processes of carcinogenesis and immune response (5). | Wild Type |
| *NFXL1* | 0.0150 | -0.75 | 28 | 62 | Transcription factor, regulation of gene expression (6). | Wild Type |
| *CORIN* | 0.0150 | -0.75 | 28 | 62 | A transmembrane serine protease, highly expressed in the heart; it participates in developmental processes and cell differentiation, particularly in the cardiovascular system (7). | Wild Type |
| *OCIAD1* | 0.0124 | -0.76 | 29 | 61 | Regulates energy metabolism and the activity of mitochondrial complex I; influences the early formation of mesodermal progenitor cells (8). | Wild Type |
| *COMMD8* | 0.0150 | -0.75 | 28 | 62 | COMMD8 is part of a complex with other COMMD proteins, CCDC22, and CRL1, which ubiquitinates κb, activating NF-κb and regulating immune response genes, cell survival, and proliferation. It also participates in copper homeostasis, sodium balance, and adaptation to hypoxia (9). | Wild Type |

**Survival Analysis WES OV**

To evaluate the prognostic significance of indel burden and indel length characteristics, overall survival (OS) was analyzed using the Kaplan-Meier method (Supplementary Figures 2 and 3). Patients were stratified into two groups based on the median value of each parameter: below the median (< Median) and equal to or above the median (≥ Median). To ensure methodological robustness, all analyses were performed independently using variants identified by three different variant-calling algorithms: Mutect2, Pindel, and VarScan2. Statistical significance of differences between survival curves was assessed using the log-rank test.

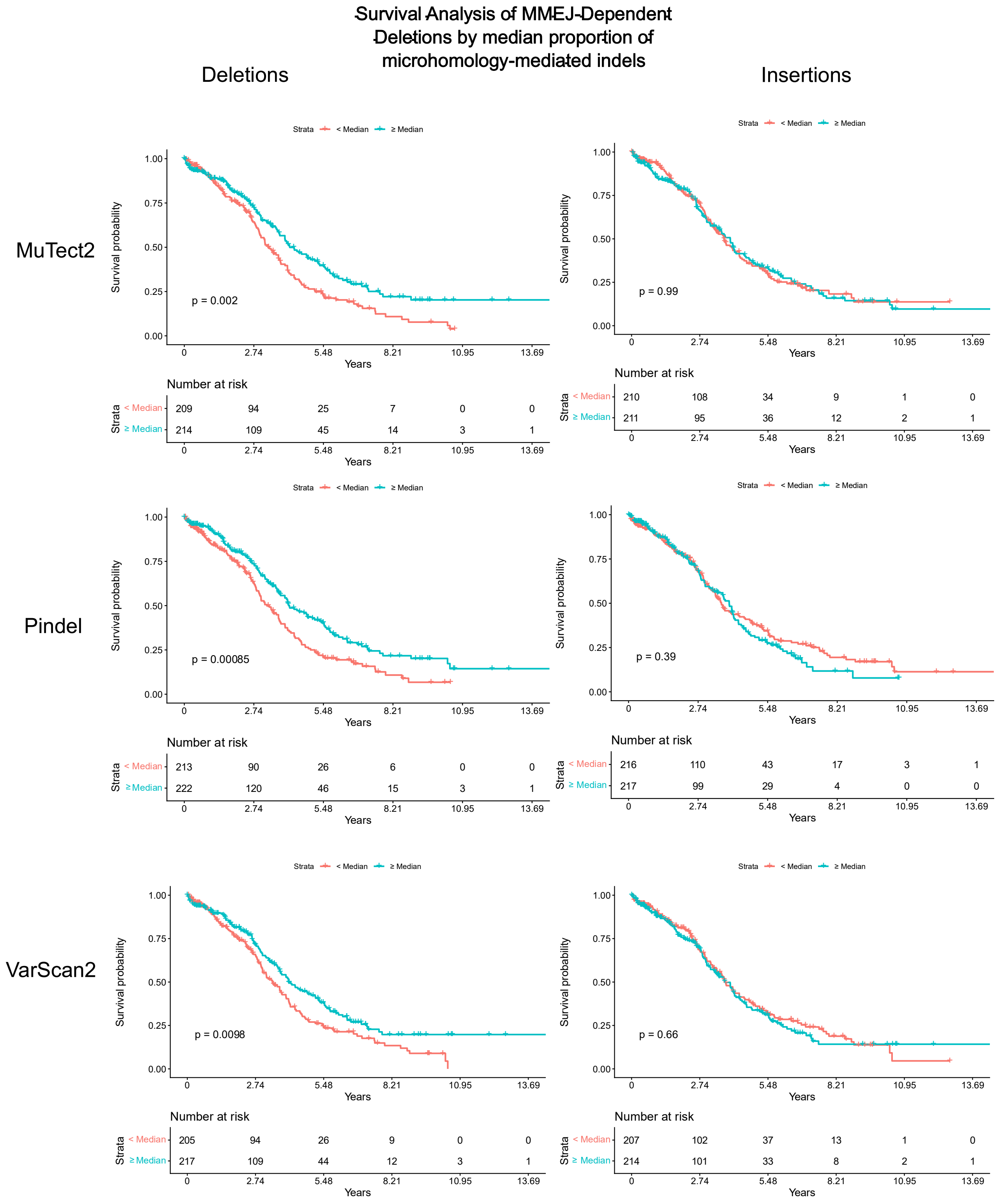


*Supplementary Figure 2 Kaplan-Meier survival analysis according to the percentage of deletions and insertions identified by the Mutect2, Pindel, and VarScan2 algorithms. The panels present survival curves where the grouping was performed analogously based on the median percentage of indel in the genome: a value below the median (red line, < Median) and a value equal to or above the median (blue line, ≥ Median). The significance of differences between groups was assessed using the log-rank test (p is given in the graphs). The lower part of the panels contains tables of the number of patients at risk (number at risk) in subsequent years of follow-up.*

Survival Analysis of Average Microhomology
Length by median median proportion of
microhomology-mediated length of indels

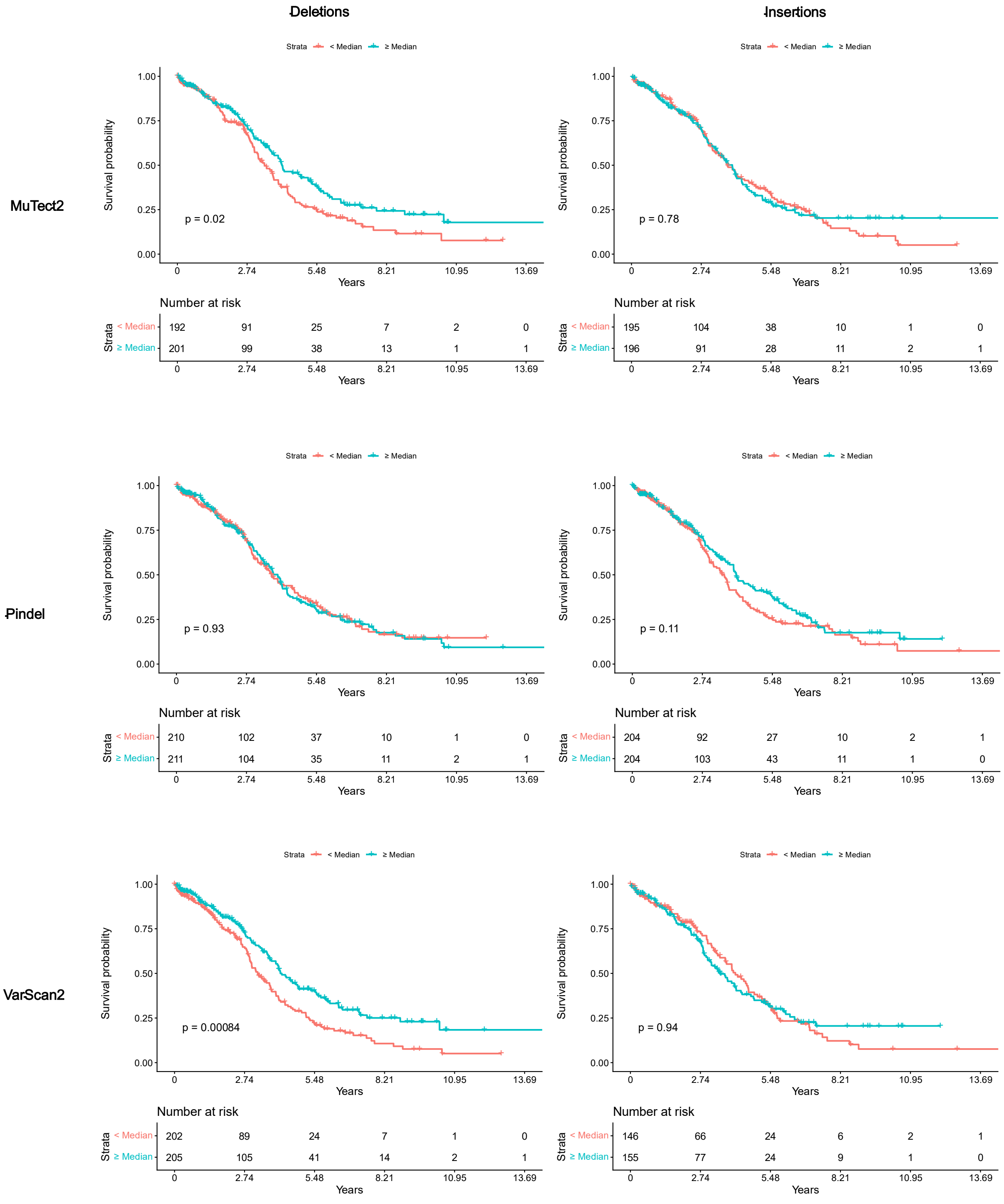


*Supplementary Figure 3 Kaplan-Meier survival analysis according to the cumulative length of deletions and insertions identified by the Mutect2, Pindel, and VarScan2 algorithms. The panels present survival curves in which patients were substratified into two groups based on the median for the cumulative length of variants: a value below the median (red line, < Median) and a value equal to or above the median (blue line, ≥ Median). Differences in overall survival were assessed using the log-rank test (p values are provided directly on the graphs). The lower part of the panels contains tables of the number of patients at risk (number at risk) in subsequent years of follow-up.*

### Technical Validation of Sequencing Methods: WGS versus WES

A comparison of WGS and WES data is presented in Supplementary Figure 4. Correlation analysis of the median indel burden measured by both methods at the cohort level demonstrated strong statistical concordance ($R = 0.89$, $p = 6.9 \times 10^{-12}$) (Supplementary Figure 4A). In contrast, the proportion of microhomology-mediated deletions in matched sample pairs showed a moderate correlation ($R = 0.59$, $p = 9.5 \times 10^{-256}$) (Supplementary Figure 5B). These findings indicate that WES provides high accuracy for estimating MMEJ burden at the population level while also highlighting its limitations in the precise quantification of microhomology-mediated deletions in individual patients.

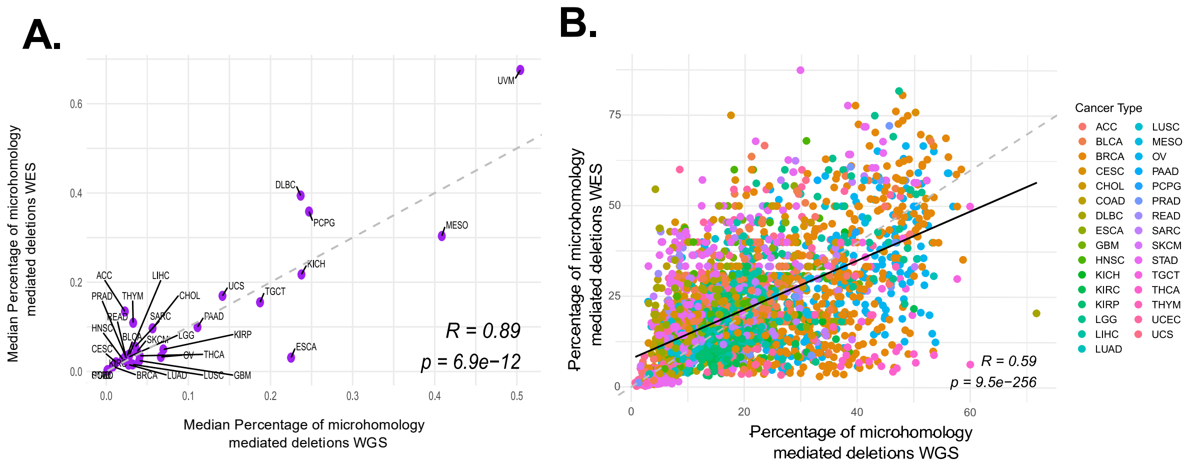


*Supplementary Figure 6 Comparison of WGS- and WES-derived estimates of microhomology-dependent deletions in matched Pan-Cancer TCGA samples. A. Scatter plot comparing the median indel burden estimated from WGS (x-axis) and WES (y-axis) data across cancer types. A strong positive correlation was observed ($R = 0.89$, $p = 6.9 \times 10^{-12}$), indicating high concordance between sequencing approaches at the cohort level. B. Scatter plot assessing concordance of the proportion of MMEJ-associated deletions between matched WGS and WES samples from TCGA. A moderate correlation was detected ($R = 0.59$, $p = 9.5 \times 10^{-256}$), suggesting reduced accuracy of WES for patient-level quantification of deletion repair signatures.*

### Impact of Specific Types of Genetic Alterations on MMEJ

Supplementary Table S2 presents the results of a generalized linear model (GLM) analysis evaluating the impact of specific types of genetic alterations, single-nucleotide variants (SNVs), copy number variations (CNVs), and structural variants (SVs), on the proportion of microhomology-mediated deletions across cancers. The odds ratio (OR) values reflect the strength and direction of the association for each alteration. OR values substantially greater than 1 indicate a markedly increased likelihood of a higher burden of microhomology-mediated deletions in the presence of these alterations. As in the previous table, statistical significance was assessed using the Benjamini-Hochberg procedure (adjusted *p*-values), and the table is supplemented with a description of the key biological functions of the analyzed genes.

*Supplementary Table 2 Genes identified by GLM analysis showing significant associations between specific mutation types and MMEJ burden. The table presents the characteristics of genes identified using a Generalized Linear Model (GLM). The analysis evaluated the impact of specific alterations (SNV, CNV, SV) on the proportion of MMEJ-dependent deletions across cancer types. Odds Ratio (OR) values indicate the magnitude of the effect, where values greater than 1 reflect an increased likelihood of a higher MMEJ mutation burden in the presence of the specified variant. The biological function of each gene is provided to highlight its potential role. Abbreviations: BH - Benjamini-Hochberg procedure for p-value adjustment; CNV - Copy Number Variation; GLM - Generalized Linear Model; SNV - Single Nucleotide Variant; SV - Structural Variant.*

| ***Gene Symbol*** | ***Mutation Type*** | ***Odds Ratio*** | ***Adjusted P-Value (BH)*** | ***Function*** |
|---|---|---|---|---|
| FDX1 | CNV | 4.48 | 3.18E-04 | A mitochondrial iron-sulfur protein that transfers electrons from NADPH to cytochrome P450 enzymes, playing a crucial role in steroid, cholesterol, and bile acid metabolism. It is essential for metabolic homeostasis, and its alterations or deletion are implicated in lipid metabolism disorders, steatohepatitis, and cancer development (10). |
| *PDE8B* | CNV | 20.61 | 4.36E-02 | PDE8B (Phosphodiesterase 8B) encodes a high-affinity, cAMP-specific phosphodiesterase that negatively regulates the cAMP/PKA signaling pathway, thereby modulating the secretion of thyroid-stimulating hormone (TSH) and thyroid hormones, as well as steroid hormone biosynthesis and insulin secretion. Common genetic variants in PDE8B significantly influence thyroid physiology and disease susceptibility, whereas pathogenic mutations are directly associated with endocrine and neurological disorders, including primary pigmented nodular adrenocortical disease (PPNAD) (11–13). |
| *FCGR1A* | SNV | 0.31 | 2.43E-10 | FCGR1A encodes CD64, a high-affinity Fc-gamma receptor transmembrane glycoprotein that plays a critical role in both innate and adaptive immune responses. In addition, FCGR1A is closely associated with interactions within the tumor microenvironment and contributes to the pathogenesis of several cancers, including cervical squamous cell carcinoma and endocervical adenocarcinoma (CESC), cholangiocarcinoma (CHOL), kidney renal clear cell carcinoma (KIRC), and skin cutaneous melanoma (SKCM) (14). |
| *ORC5* | SNV | 0.41 | 2.20E-03 | The origin recognition complex (ORC), including its ORC2 and ORC5 subunits, is an essential protein complex responsible for binding replication origins and initiating DNA replication in eukaryotic cells. However, studies in human cancer cells have demonstrated that these cells can survive and initiate DNA replication normally even in the complete absence of ORC2 or ORC5 expression, suggesting the existence of alternative mechanisms that enable DNA replication independently of the fully assembled ORC complex (15). |
| *OSR1* | SNV | 0.36 | 7.26E-07 | OXSR1 (oxidative stress-responsive kinase 1), also known as OSR1, is a member of the Ste20 |

| | | | | |
|---|---|---|---|---|
| | | | | family of serine/threonine kinases that is widely expressed across multiple tissues, including the heart, liver, kidneys, and lungs. Its primary functions are the regulation of epithelial ion transport and the maintenance of cellular volume homeostasis (16). |
| *POU2F3* | SNV | 0.39 | 1.84E-03 | POU2F3 is the master transcriptional regulator that specifies tuft cell identity. In addition, it defines the SCLC-P molecular subtype of small cell lung cancer, which is characterized by a complete lack of neuroendocrine marker expression (17). |
| *PSMC2* | SNV | 0.40 | 1.96E-06 | PSMC2 is a key component of the 26S proteasome, which plays an important role in promoting the development of various cancers, including highly aggressive gliomas. The expression of this gene in tumor tissues has prognostic value for patients and is closely associated with DNA methylation, cell cycle regulation, and immune cell infiltration (18). |
| *SPDYE3* | SNV | 0.34 | 6.24E-03 | SPDYE3, a member of the Speedy/Ringo gene family, is a regulator of the cell cycle and DNA damage response that functions through binding to and activating cyclin-dependent kinases (CDKs). This gene is significantly overexpressed in multiple cancer tissues, including lung squamous cell carcinoma (LUSC), suggesting its potential clinical relevance and mechanistic involvement in promoting cellular proliferation (19). |
| *TBCEL* | SNV | 2.12 | 2.97E-02 | TBCEL encodes a protein involved in microtubule disassembly, which disrupts tubulin heterodimers, prevents their polymerization, and targets them for proteasomal degradation. This gene is highly expressed in the testes, where it is essential for the proper progression of spermatid individualization by initiating the required disassembly of the microtubule cytoskeleton (20). |
| *TNFRSF10B* | SV | 3.12 | 3.39E-02 | TNFRSF10B encodes the death receptor DR5, and its transcription, regulated by the ERV9-LTR element, exerts a strong pro-apoptotic function that prevents the survival of damaged germ cells. Pharmacological induction of TNFRSF10B expression, for example through the use of HDAC inhibitors, leads to rapid tumor cell death, representing a potential mechanism for restoring tumor suppression in testicular cancer (21). |

14. Frontiers | FCGR1A Serves as a Novel Biomarker and Correlates With Immune Infiltration in Four Cancer Types [Internet]. [cited 2026 Aug 4]. Available from:

https://www.frontiersin.org/journals/molecular-biosciences/articles/10.3389/fmolb.2020.581615/full

15. Shibata E, Dutta A. A human cancer cell line initiates DNA replication normally in the absence of ORC5 and ORC2 proteins. J Biol Chem. 2020;295:16949–59.

16. Gagnon KB, Delpire E. Molecular Physiology of SPAK and OSR1: Two Ste20-Related Protein Kinases Regulating Ion Transport. Physiol Rev. 2012;92:1577–617.

17. Szczepanski AP, Tsuboyama N, Watanabe J, Hashizume R, Zhao Z, Wang L. POU2AF2/C11orf53 functions as a coactivator of POU2F3 by maintaining chromatin accessibility and enhancer activity. Sci Adv. 2022;8:eabq2403.

18. Wang Y, Zhang S, Zhao Z, Jin Q, Wang Z, Song Z, et al. PSMC2 promotes glioma progression by regulating immune microenvironment and PI3K/AKT/mTOR pathway. Immunobiology. 2024;229:152802.

19. Shao Z, Ye J, Wu Y, Chen Y, Wang R, Fang S, et al. SPDYE3 promotes cell cycle and LUSC progression by regulating the CDC25C/CDK1 pathway. Int J Biochem Cell Biol. 2025;186:106825.

20. Fabrizio JJ, Rollins J, Bazinet CW, Wegener S, Koziy I, Daniel R, et al. Tubulin-binding cofactor E-like (TBCEL), the protein product of the mulet gene, is required in the germline for the regulation of inter-flagellar microtubule dynamics during spermatid individualization. Biol Open. 2020;9:bio049080.

21. Beyer U, Krönung SK, Leha A, Walter L, Dobbelstein M. Comprehensive identification of genes driven by ERV9-LTRs reveals TNFRSF10B as a re-activatable mediator of testicular cancer cell death. Cell Death Differ. Nature Publishing Group; 2016;23:64–75.
